# Bioindicator-based mapping of peatland potential in the Western Mediterranean-Atlantic Realm

**DOI:** 10.1101/2025.03.21.644445

**Authors:** Miguel Geraldes, Raquel Fernandes, Maurício Santos, César Capinha

**Author notes:** **Corresponding author:** (R. Fernandes).

## Abstract

Ecosystems with a naturally accumulated peat layer at the surface and wetlands with or without a peat layer dominated by a vegetation that may produce peat (hereafter ‘peatlands’ for simplicity) are critical assets in global change due to their large carbon content storage capacity and essential ecosystem services. A Global Peatland Assessment was initiated during the UNFCCC COP22; however, many countries still lack accurate peatland maps. This is particularly concerning in the Western Mediterranean, where these ecosystems are frequently overlooked or misclassified. To help address this issue, we developed predictive distribution models for vascular plant and bryophyte species typical of peatland-type systems in the Western Mediterranean. We identified 28 ‘indicator’ mire-typical plant taxa and created a database using filtered data from six different data sources, including our own field surveys. Thirteen high-resolution spatial predictor variables, covering topographic, soil, and hydroclimatic factors, were selected to reflect potential relationships with species presence. For each species, a potential distribution model was built using MaxEnt software. The resulting potential-distribution maps were cross-validated, and models with an AUC score of ≥0.75 were classified into suitable or unsuitable areas. The species richness map aligned well with areas known to support peatlands. The indicator species-based mapping also suggests a broad potential for yet unrecorded peatlands in the Western Mediterranean, including small areas, particularly along the western and southwestern sandy coastlines and in the southern mountains.

## 1. Introduction

The global importance of peatlands for Humankind has become a hot topic since they provide major carbon storage and sequestration services, among other services, such as regulation of water quality, biodiversity support, energy and material provision and cultural activities (Joosten & Clarke, 2002). Peatlands gained significant global recognition at the United Nations Framework Convention on Climate Change (UNFCCC) COP22 held in Marrakesh, Morocco, in 2016. This marked a pivotal moment in the global environmental agenda as the importance of peatlands in climate change mitigation and biodiversity conservation was emphasized. At COP22, discussions stressed the need for the protection and sustainable management of peatlands to achieve the goals of the Paris Agreement (COP21) and combat climate change effectively. This was a significant step in elevating the conservation of peatlands onto the international climate agenda (Tanneberger et al., 2021).

To prevent the loss of peatland areas, it is essential first to identify where they occur. However, comprehensive distribution data for peatland ecosystems remain incomplete across many regions worldwide (Minasny et al., 2019). The Western Mediterranean (WM) is one such region where significant gaps in peatland mapping persist. In the Iberian Peninsula, peatlands are predominantly described in the northern and northwestern mountain ranges (e.g., Cantabrian Mountains; Chico et al., 2019), where high precipitation and glacial or periglacial landforms have fostered water retention and peat accumulation through processes like paludification or the terrestrialization of basins (Martínez-Cortizas et al., 2000). Other areas, namely the Toledo Mountains in central Iberia (López-Sáez et al., 2014), and the Gredos range in the Iberian Central System (López-Sáez et al., 2023a), have been largely surveyed to characterise and describe the current conditions of peatlands here located.

However, substantial knowledge gaps remain particularly in the southern Iberian Peninsula and northern Maghreb, and this underrepresentation likely stems from the perceived lower prevalence of peatlands. In these areas, peatlands are often described as small and fragmented environments with low peat accumulation rates (Pontevedra-Pombal et al., 2017), owing, to a large extent, to dry and warm climate conditions that lead to lower water availability (Tanneberger, Moen, et al., 2021). However, significant peat accumulation also occurs in some coastal and sublittoral wetlands of the western, southwestern, and southern regions (Mateus et al., 2017). These include interdune slacks, fluvial endorheic mires on blocked fluvial catchments, and back-swamp fluvial mires — low-water flow environments where unique conditions have allowed peat formation and thick accumulation even under a warmer climate (Pontevedra-Pombal et al., 2017). Peat deposits often emerge as a subtle landscape mosaic in these areas, making them less extensive than their boreal counterparts but notable for their higher floral diversity (Joosten et al., 2017). Similarly, knowledge about peatlands in northwestern Africa is fragmented, with studies highlighting significant peat deposits in northern Morocco (Muller et al., 2011) and northeast Algeria-northern Tunisia (Muller et al., 2010).

Although, in the last decades, these peatlands have been experiencing strong pressure from multiple human activities (Heras-Pérez et al., 2017; Mateus et al., 2017), leading to the reduction of their extent or total disappearance (Tanneberger, Moen, et al., 2021), several peatland areas persisted in this region (Fernandes et al., 2024), and improved mapping efforts are necessary for accurately understanding their current distribution.

Additionally, accurately mapping is crucial to identifying and prioritizing conservation and restoration areas, preserving their critical ecological roles, including supporting a diverse array of species of conservation concern (Neto et al., 2021), carbon storage and climate regulation (Tanneberger, Appulo, et al., 2021), and assess the possible impacts of several anthropogenic pressures (Fernandes et al., 2024).

There are multiple approaches to digital mapping peatlands (Minasny et al., 2024). Vegetation cover, particularly peat indicator species, has been used as a proxy to assess the potential distribution of peatlands through species distribution models, mainly in temperate and boreal regions, where peatlands are mainly dominated by *Sphagnum* mosses (Oke & Hager, 2017; Campbell et al., 2021). Species distribution models (SDMs), which use species occurrence records and associated environmental conditions (Pearson, 2010), allow the prediction of the distribution of mire-typical indicator species (Hammerich, Damman, et al., 2022), enabling identifying locations where environmental conditions may be suitable for peatlands to occur (Cong et al., 2020). In particular, estimates of co-occurrence among these species, represented as species richness maps, offer a broader, cross-taxa perspective on where peat-forming environments are most likely to occur (Saarimaa et al., 2019) or predict shifts in their location (Ma et al., 2022). This approach minimizes reliance on the ecological requirements of individual species and instead highlights areas of shared suitability across multiple taxa (Oke & Hager, 2017). Moreover, richness-indicative models provide a robust baseline for guiding field surveys and can strategically inform mapping efforts in regions where peatland data are sparse or outdated (Melton et al., 2022).

Here, we aim to support future efforts of mapping peatlands in the Iberian Peninsula and NW Africa by producing mire-typical species (Hammerich, Damman, et al., 2022) distribution maps. We use a comprehensive dataset of species occurrence records derived from multiple sources, coupled with high-resolution environmental predictors representing topographic, soil, and climatic variables. By applying species distribution models, we aim to predict the potential distribution of peatland indicator species and identify regions with high species richness that align with peat-forming environments. This approach not only helps to locate overlooked peatland areas but also provides an initial baseline for accurate ground-truthing and conservation planning, prioritizing efforts in regions of high vulnerability and ecological significance.

## 2. Materials and Methods

### 2.1 Study area

The Iberian Peninsula (Portugal and Spain) and the north coast of Africa (Morocco and Algeria), the study area, are located in the Western Mediterranean (approximately between 33°25′ N and 43°47′ N and 9°00′ W and 8°25′ E), being surrounded by the Mediterranean Sea, the Cantabrian Sea, and the Atlantic Ocean (fig.1). While the Mediterranean climate is dominant in the core and south of the Iberia Peninsula and along the north coast of Africa, the temperate oceanic climate is dominant in the north of the Peninsula and has important influence along the western coastline (Kottek et al., 2006). The Iberian Peninsula is mainly covered by plains and surrounded by mountain chains (Pontevedra-Pombal et al., 2017). In North Africa, the Mediterranean coasts of Morocco and Algeria are mostly mountainous albeit coastal sedimentary plains dominate along the NW Moroccan Atlantic coast (Britton & Crivelli, 1993). The study area is located in a border zone between the Eurosiberian and Mediterranean floral regions, influencing the distribution and characteristics of the different types of peatlands (Pontevedra-Pombal et al., 2017). The Eurosiberian region is the most suitable region for peatland development (Heras-Pérez et al., 2017), while Mediterranean peatlands are considered groundwater-dependent (particularly during summer), small, scattered, and shaped by the local environment (Santoni et al., 2021). Peatlands in the Iberian Peninsula and northern Africa share ecological and biogeographical similarities, forming a connected Mediterranean peatland framework, despite some genetic differentiation (Geraldes et al., 2014, Neto et al., 2019).

**Figure 1.**
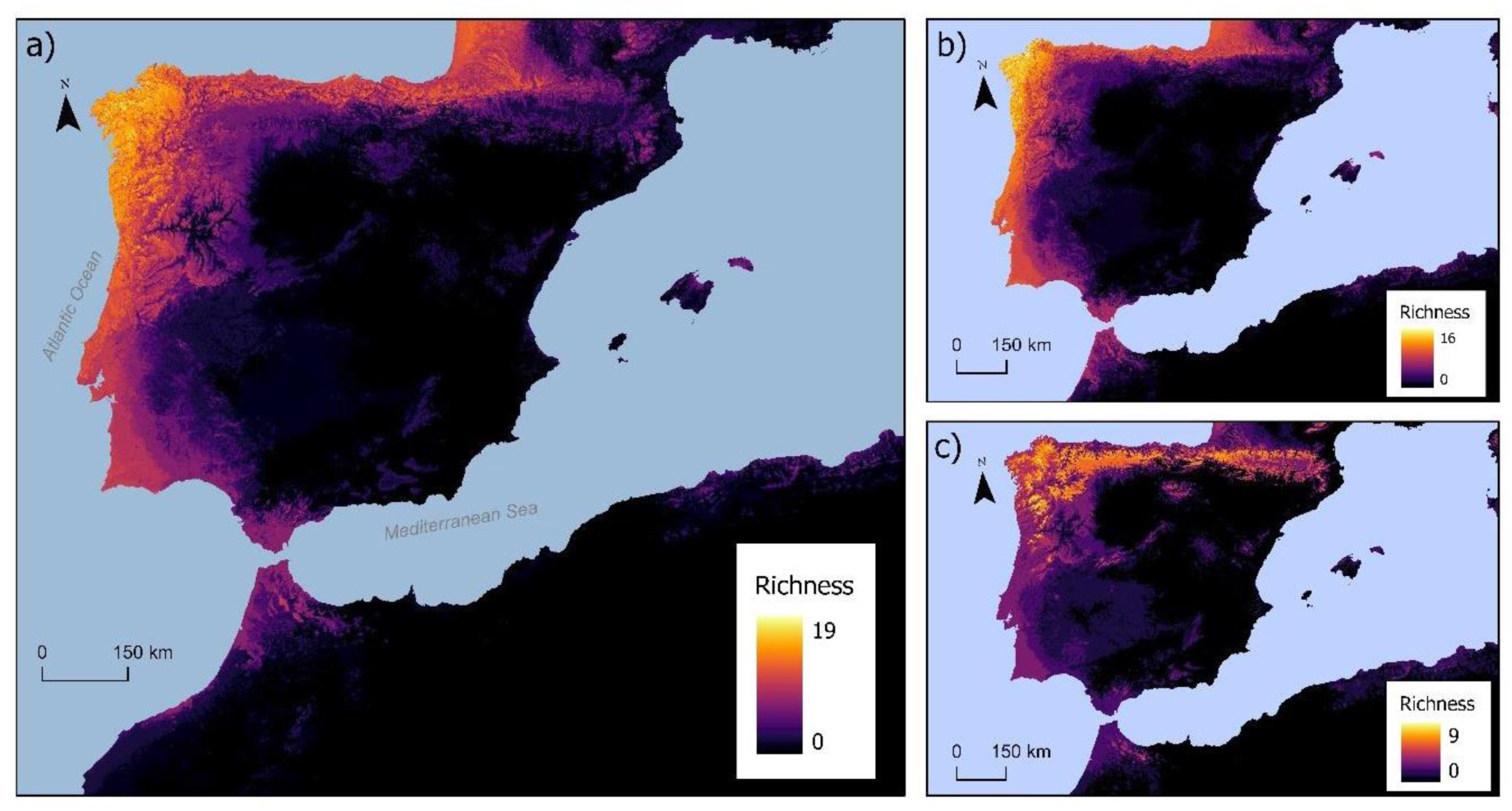
Predicted richness of plant taxa associated with mires, peatlands and other potential organic soil-forming environments in the Western Mediterranean Realm. Richness maps are presented for all taxa (a), for taxa found in lower-altitude environments (b), and for taxa found in higher-altitude environments (c).

### 2.2. Mire-typical indicator species

We selected a total of 28 mire-typical indicator taxa, comprising 25 vascular plants (including two subspecies) and 2 moss species, known to be associated with peatlands and other organic soil-forming environments across the subregions of the Iberian Peninsula and Northwestern Africa, after Castroviejo et al. (1986), Séneca (2003), Romo (2009), Muller et al. (2010, 2011), and Neto et al. (2019, 2021) (Table 1). While any selection of indicator taxa has inherent limitations—varying degrees of specialization to the ecosystems of interest and uneven representation across the full gradient of environmental conditions—our approach prioritized capturing the diversity of peatland conditions found in the Iberian and Northwest African contexts by including an array of genera of mire-typical plants that collectively reflect the distinct environmental conditions of the study area. This included balancing mire-typical species of high-altitude environments with those associated with low-altitude ecosystems, such as the southern Iberian plains and coastal regions (Table 1). To map the distribution of selected taxa, we compiled occurrence records from multiple reference databases, including Anthos (anthos.es), Flora-On (flora-on.pt), SIVIM (sivim.info), and GBIF (gbif.org), supplemented by data from additional literature and our own field records. Importantly, the distribution of each taxon was mapped across the entire extent of Europe and North Africa, rather than being limited to the boundaries of the study area. This broader geographical scope allowed us to more accurately capture the range of environmental conditions that define each species’ ecological niche, and mitigate niche truncation, where models based on limited data fail to fully capture a species’ ecological niche, leading to biased or incomplete distribution predictions (Chevalier et al., 2022)

**Table 1.** Taxa used as mire-typical indicators in the Iberian and Northwest-African region. The species list reflects the diverse environmental and habitat conditions, with regional notorious prevalence of fens over moss-dominated bogs, in which vascular plants (Va) and bryophytes (Br) are found within likely peat-forming areas in the study area. It specifically distinguishes taxa commonly associated with higher-altitude (A) conditions from those found in lower-altitude or even coastal environments (L). Additionally, the number of distribution records used for model fitting is provided, along with the average area under the curve (AUC) across replicate models and its standard deviation. AUC values and standard deviation for taxa surpassing the threshold of performance defined for inclusion in richness maps are shown in bold.

| Species | Altitudinal association | Group | Number of records | Average AUC | AUC Standard deviation |
| --- | --- | --- | --- | --- | --- |
| *Agrostis juressi* | A; L | VA | 9 | **0.98** | **0.02** |
| *Anagallis tenella* | L | VA | 3,186 | **0.88** | **0.002** |
| *Arnica montana* subsp. *atlantica* | A | VA | 52 | **0.99** | **0.004** |
| *Arnica montana* subsp. *montana* | A | VA | 55 | **0.97** | **0.019** |
| *Carex demissa* | L | VA | 20,715 | 0.68 | 0.002 |
| *Cirsium palustre* | A; L | VA | 133,521 | 0.54 | 0.002 |
| *Cladium mariscus* | L | VA | 4,060 | **0.85** | **0.007** |
| *Drosera intermedia* | L | VA | 6,774 | **0.81** | **0.003** |
| *Drosera rotundifolia* | A | VA | 3,387 | 0.61 | 0.003 |
| *Erica ciliaris* | L | VA | 2,140 | **0.92** | **0.001** |
| *Erica tetralix* | A | VA | 42,250 | 0.61 | 0.001 |
| *Genista ancistrocarpa* | L | VA | 151 | **0.99** | **0.001** |
| *Genista anglica* | A | VA | 8,927 | **0.79** | **0.002** |
| *Genista berberidea* | L | VA | 35 | **0.99** | **0.001** |
| *Gentiana pneumonanthe* | A; L | VA | 7,460 | **0.78** | **0.003** |
| *Juncus bulbosus* | L | VA | 29,061 | 0.64 | 0.004 |
| *Lobelia urens* | A; L | VA | 2,209 | **0.92** | **0.003** |
| *Molinia caerulea* | A; L | VA | 103,316 | 0.55 | 0.003 |
| *Myrica gale* | L | VA | 25,282 | 0.66 | 0.003 |
| *Pinguicula lusitanica* | L | VA | 2,734 | **0.93** | **0.000** |
| *Potentilla erecta* | A; L | VA | 128,239 | 0.54 | 0.001 |
| *Rhynchospora alba* | A | VA | 10,507 | **0.75** | **0.003** |
| *Rhynchospora modesti-lucennoi* | L | VA | 33 | **0.99** | **0.002** |
| *Sphagnum auriculatum* | A; L | BR | 286 | **0.98** | **0.003** |
| *Sphagnum subnitens* | A; L | BR | 9,053 | **0.78** | **0.006** |
| *Thelypteris palustris* | L | VA | 8,194 | **0.77** | **0.005** |
| *Utricularia australis* | L | VA | 4,111 | **0.83** | **0.008** |
| *Utricularia gibba* | L | VA | 16 | **0.81** | **0.138** |

To ensure high-quality data, the occurrence dataset was processed and cleaned, including coordinate standardization (converting UTM and degrees/minutes/seconds to decimal degrees) and thorough data validation using the CoordinateCleaner package for R (Zizka et al., 2019) to remove duplicates, errors, and geographically implausible records.

### 2.3 Predictor variables

We used 13 environmental predictor variables (Table 2) based on their potential to determine the distribution of the selected indicator taxa. These variables can be broadly grouped into three main categories: topographic, soil, and bioclimatic variables. Topographic variables include elevation, slope, aspect, flow accumulation, and topographical wetness index and aim at representing topographic variation, which is a major factor controlling the development of organic soils in areas with a permanent water surplus and provide key insights into hydrological processes influencing peatland formation (Barthelmes et al., 2015). Soil variables were obtained from SoilGrids (Poggio et al., 2021), including bulk density, soil organic carbon (SOC) stock, pH, and soil texture. To account for hydro-climatic influences on WM peatlands, we compiled a set of climatic variables. These variables, focusing on extreme or limiting conditions of temperature and precipitation, were obtained from the CHELSA Bioclim dataset for the reference period 1979–2013 (Karger et al., 2017). They are particularly relevant for understanding seasonal recharge mechanisms in Mediterranean peatlands (Santoni et al., 2021) and for species distribution modelling in these climatically variable regions. By integrating these topographic, soil, and bioclimatic variables, this study leverages a comprehensive set of predictors to model peatland distribution and the environmental niches of key indicator species.

**Table 2.**
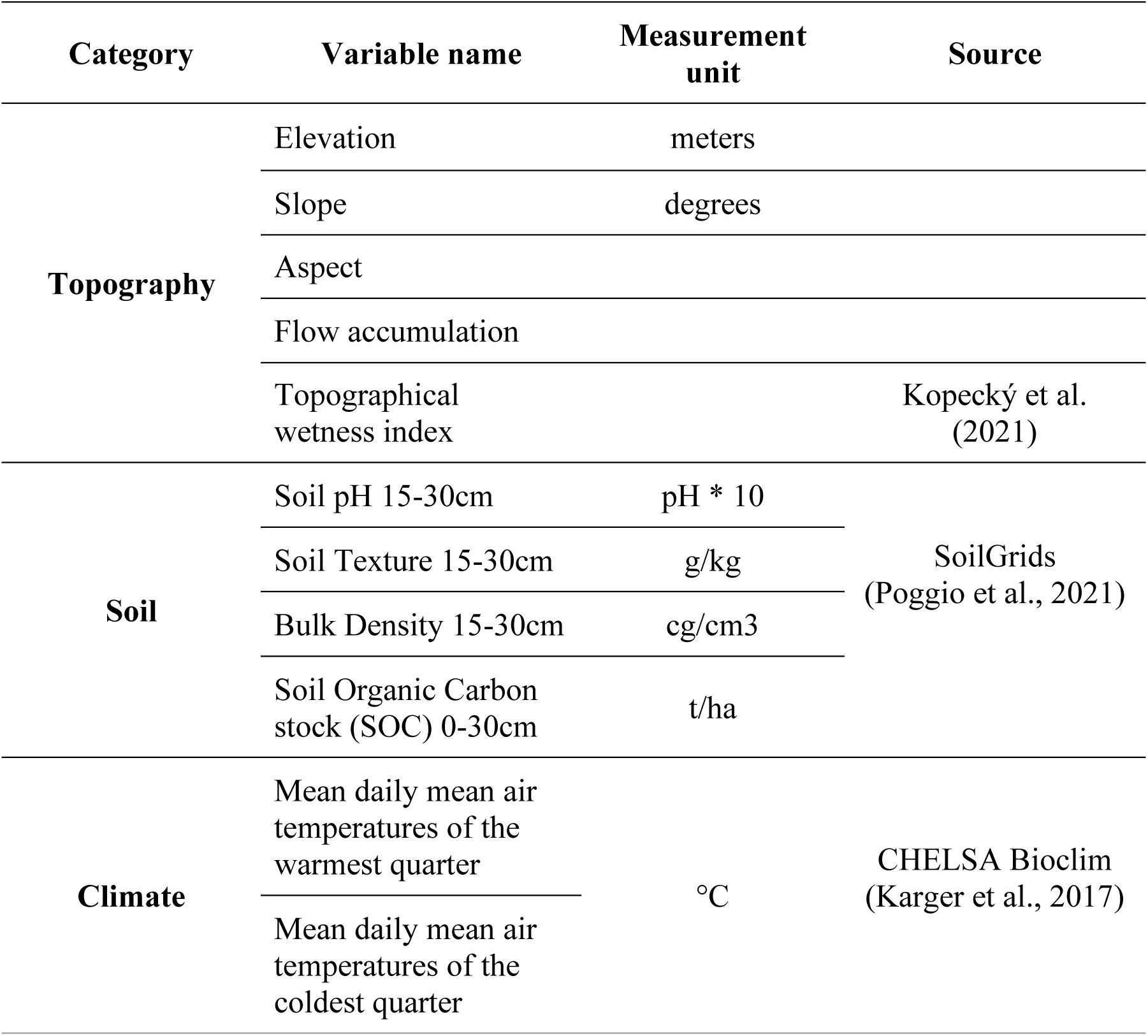
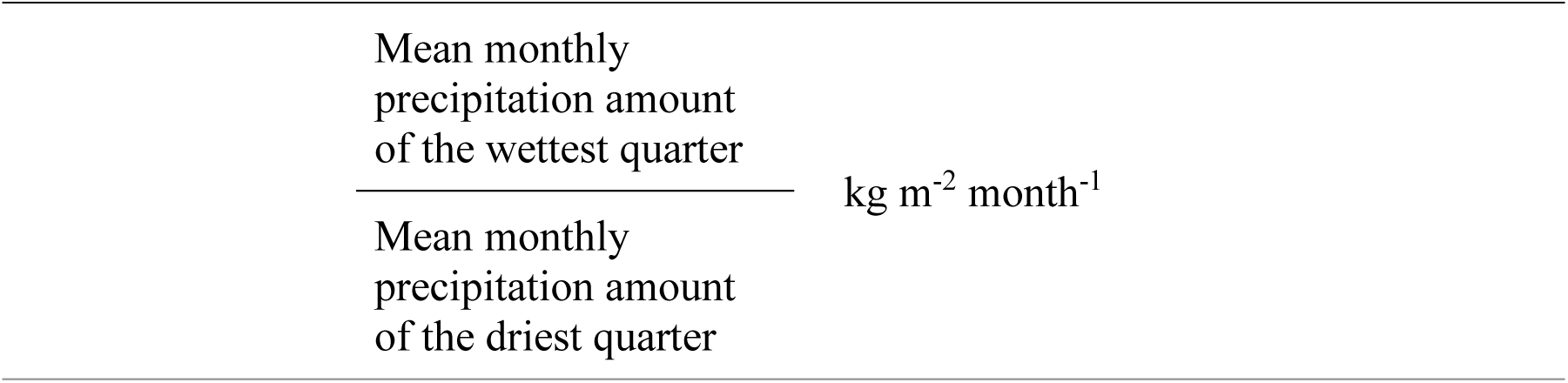
Predictor variables considered for modelling the potential distribution of indicator taxa.

### 2.4 Modelling of the potential distribution of indicator taxa

To predict the potential distribution of selected peat indicator taxa, we employed the widely used Maximum Entropy (MaxEnt) modeling software, which is an open-source, Java-based application that uses a machine-learning algorithm grounded in the principle of maximum entropy (Phillips & Dudík, 2007; Pearson, 2010). It offers several advantageous properties, including insensitivity to collinearity among predictors, robustness even with a low number of species occurrence records, and the ability to generate comprehensive diagnostic reports to assess model performance (Pearson, 2010). Moreover, MaxEnt identifies geographic regions with suitable environmental conditions for species by analysing species occurrence data alongside background environmental information (Elith et al., 2011) To implement the models, we used 10,000 background points to represent available environmental conditions and cloglog as the output format, where predicted environmental suitability values vary between 0 and 1. The same background environmental data was used as a proxy of the potential distribution of the peat indicator species, regardless of their mire-typical ecological niche conditions.

We measured the performance of models for each taxa using the area under the curve (AUC) of receiver operating characteristic plots (Fielding & Bell, 1997). To perform this assessment, four replicate models were developed for each taxon. In each, 75% of occurrence data were randomly selected for model training, while the remaining 25% were reserved for validation (Pearson, 2010; Ma et al., 2022). A cross-replicate average AUC value of ≥0.75 was considered indicative of a model with sufficient accuracy for representing the species potential distribution (Elith & Burgman, 2002; Pearson, 2010).

For each replicate of species with an average AUC ≥0.75, we obtained the maximum test sensitivity plus specificity (maxTSPS) logistic threshold, following (Liu et al., 2005). This threshold represents the predicted probability value that optimally balances the model’s ability to accurately predict species presence (sensitivity) and absence (specificity). This approach is particularly recommended for models employing pseudo-absence or background data, as it ensures an effective trade-off between minimizing false positives and false negatives, thereby enhancing the reliability of derived binary classifications (Elith et al., 2011).

Using the mean maxTSPS threshold, we reclassified the average MaxEnt replicate predictions for each species into binary presence-absence maps, delineating areas of predicted occurrence for each regionally mire-typical indicator taxon. To evaluate potential species richness within the study area, these binary maps were summed, creating richness maps that represent areas of potential peatland indicator diversity. We generated richness maps for the full suite of indicator taxa, as well as for taxa occurring in lower-altitude or even coastal environments and those occurring in higher-altitude conditions, providing a nuanced view of biodiversity patterns across environmental gradients of the study area.

## 3. Results

A total of 20 (out of 28) taxa achieved cross-validation performance values above the AUC threshold defined for inclusion in richness maps (i.e., AUC ≥ 0.75; Table 1). Half of these species achieved AUC values above 0.9 (n =10): *Agrostis juressi, Anagallis tenella, Arnica montana* (both subspecies), *Erica ciliaris, Genista ancistrocarpa, Genista berberidea, Lobelia urens, Pinguicula lusitanica, Rhynchospora modesti-lucennoi, and Sphagnum auriculatum*. These models also exhibit low standard deviations (<0.01), indicating stable predictions across replicates. A few species with borderline AUC values between 0.75 and 0.79, are *Gentiana pneumonanthe*, *Genista anglica, Rhynchospora alba, Sphagnum subnitens,* and *Thelypteris palustris*. Conversely, species with AUC < 0.75, such as *Carex demissa, Cladium mariscus* (low-performing replicate), *Drosera rotundifolia, Erica tetralix, Juncus bulbosus, Molinia caerulea, Myrica gale*, and *Potentilla erecta*, failed to achieve the required performance and were excluded from further analysis.

The predicted richness of indicator species exhibits distinct spatial patterns across the Iberian Peninsula and northern Morocco (Fig. 1). The aggregate richness of all indicator species (Fig. 1a) reveals high concentrations of richness in the northern regions of the Iberian Peninsula, particularly in the Pyrenees, Basque-Cantabrian Mountain ranges, and along the northern Atlantic coastline, with a peak in Galicia and Northwest Portugal. Richness decreases significantly toward the southern half of the Iberian Peninsula and northern Morocco and the northernmost Algerian-Tunisian border region, where large areas show minimal or no predicted richness. These patterns suggest that regions with higher precipitation, cooler temperatures, and favorable hydrological conditions serve as hotspots for mire-typical indicator species.

For species able to occur in low-altitude fen-dominated areas (Fig. 1b), richness is highest along the western coastal areas of Portugal and Galicia, extending into the Atlantic coastline. The central and southern regions of the Iberian Peninsula show moderate richness, while northern Morocco and the northernmost Algerian-Tunisian border region exhibits slightly higher richness compared to the high-altitude species map, particularly in coastal or lowland areas. These patterns reflect not only the adaptability of low-altitude taxa to environments with lower elevations and more localized or seasonal (even intermittent) water availability but also the relevance of some southern medium-altitude ridges near the Strait of Gibraltar for such taxa (e.g. Cadiz mountains, Western Rif) as well as extensive sandy coastal deposits with permanent groundwater supplies where those plants can thrive.

When focusing on high-altitude indicator species (Fig. 1c), predicted richness is highest in Spanish mountainous regions such as the Pyrenees, the Cantabrian range, and parts of the northern area, namely in León Mountains and Galicia’s ‘sierras orientales’. In Portugal, the highest richness was observed mainly in the Estrela Mountain and parts of the northern mountains (e.g. Alvão, Cabreira, Gerês, Peneda, Montesinho) with some connectivity to the Spanish mountain systems. In contrast, much of the central and southern Iberian Peninsula exhibits little to no richness, with only sparse occurrences in isolated high-altitude areas such as the Malcata-Gata, Francia, Béjar, Gredos, Guadarrama, and Ayllón Mountains in the Central Iberian System, the northern part of the Iberian System, and the Sierra Nevada. Moderate richness values persist in southern coastal areas of Portugal, sustained by taxa that can thrive across both high- and low-altitude environments. Northern Morocco shows moderate to high richness associated to Western Rif mountains, but low to no richness outside this region. These patterns align with the ecological requirements of high-altitude species, which rely on the cooler and wetter conditions typically found in mountainous environments.

When comparing the three maps, the aggregate richness map (Fig. 1a) appears strongly influenced by contributions from high-altitude taxa, as seen in the prominence of northern mountainous regions. However, low-altitude taxa contribute significantly to richness in coastal and lowland areas, particularly in southwestern Iberia and northern Morocco, showing even some ecological amplitude to climb some of the local medium-altitudinal ranges and colonize local nestled mires and peatlands. High-altitude taxa (Fig. 1c) are strongly localized to northern mountainous regions and are largely absent from lower and southern latitudes, whereas low-altitude taxa (Fig. 1b) exhibit a broader distribution that extends into southern and coastal areas.

## 4. Discussion

Our results revealed distinct spatial patterns of predicted species richness across the Iberian Peninsula, northern Morocco, and the northernmost Algerian-Tunisian border region, highlighting significant hotspots of richness in northern mountainous regions for high-altitude taxa and moderate richness values in southern coastal areas of Portugal and lowland regions of northern Morocco. Though these findings align to some extent with known peatland distributions, we also provided new information on areas with high peat-forming potential, particularly in less-studied regions, including many tiny ecosystems. One of these regions is the Portuguese Atlantic coast, especially in the central and south coastal areas. Here peatlands are known whether to be rarer promoted by springs but with little peat accumulation due to warmer and drier conditions (Tanneberger, Moen, et al., 2021), or, in coastal mires, they can sometimes store a thick peat layer and their presence is associated with specific soil conditions - high water storage coefficients that compensate for the large evapotranspiration losses in summer (Mateus et al., 2017). Yet, our results showed that, here, selected areas could be the target of fieldwork to identify peatland locations that are not identified yet.

By using a bio-indicator approach that uses highly specialized plants with great fidelity to these habitats (Geraldes, et al., 2014, Neto, et al., 2021), the predicted richness maps align with established knowledge of potential peat-forming ecosystems location in WM (Heras-Pérez et al., 2010, 2017; López-Sáez et al., 2014, 2023b; Mateus et al., 2017; Pontevedra-Pombal et al., 2017; Chico et al., 2019). High-altitude species richness was strongly associated with environments where cooler temperatures, higher precipitation, and mountainous topography are dominant, as observed in the northern regions, from the Basque - Cantabrian and Galician-Asturian areas through northern Portugal. This can be partially justified by their location in the Eurosiberian floral region, which is influenced by a humid climate moderated by the ocean and characterized by mild to cold winters, without a distinctly dry season (Pontevedra-Pombal et al., 2017) In the Eastern Pyrenees, peatlands are considered more localised and uncommon, when compared to the Basque-Cantabrian mountains, due to drier climate conditions (Heras-Pérez et al., 2017). However, in Western Pyrenees, the highest species richness can be associated with the Atlantic influence (Heras-Pérez et al., 2010).

Our findings provide critical insights into the ecological conditions that shape the distribution of peatland and organic soil-forming in lowland environments in the WM. Moderate richness values were observed in central and southern coastal areas of Portugal, driven by taxa capable of thriving in low-altitude conditions. Previous knowledge showed that peat formation in coastal and lowland regions is observed in poor drainage basins, with permanent water saturation and low water flow, even in warmer climates (Mateus et al., 2017). Additionally, low-altitude species richness in northern Morocco, northwestern Algeria and northern Tunisia reflects the adaptability of certain taxa to environments with intermittent or localized water availability, emphasizing the need to further explore and validate these areas as potential peat-forming ecosystems or ecosystems with accumulation of peat in the surface.

This study contributes methodologically by leveraging SDMs, specifically MaxEnt, to predict potential peatland distributions in a region where direct mapping remains sparse. The integration of high-resolution environmental predictors, including topographic, soil, and climatic variables, proved effective in modelling the ecological niches of peatland indicator species. Peatland mapping efforts based on bioindicator-base SDMs in other regions, particularly in boreal and temperate zones, showed their role in land-use planning, managing help considering different conservation challenges, and assessing possible changes in distribution patterns due to climate change scenarios (Oke & Hager, 2017; Saarimaa et al., 2019; Campbell et al., 2021). However, the distinct ecological and climatic conditions of the WM necessitate tailored approaches. Unlike boreal peatlands, where species such as *Sphagnum* mosses dominate (Rydin et al., 2006), Mediterranean peatlands exhibit a more diverse - namely vascular - floral composition (Joosten et al., 2017), requiring a broader set of mire-typical indicator taxa to capture the complexity of peat-rich systems.

By including both high- and low-altitude taxa, our richness maps capture a broader range of environmental gradients, reducing reliance on single-species ecological requirements and improving predictive reliability.

The use of a cross-taxa approach, particularly the richness-indicative modelling, provides a baseline for future ground-truthing surveys. This approach ensures that predictions reflect shared suitability conditions across taxa, making them more generalizable for identifying peat-forming environments. Mapping species distributions across the broader geographical extent of Europe and North Africa helped mitigate niche truncation, ensuring more accurate representations of ecological niches (Chevalier et al., 2022). Additionally, the methods and insights from this study could be extended to other underrepresented regions within the Mediterranean or globally. For example, similar bioindicator-based approaches could be applied to map peatlands in other semi-arid or coastal environments, contributing to global peatland mapping initiatives such as the Global Peatland Assessment (United Nations Environment Programme, 2022).

Our results carry relevant implications for conservation and management efforts. Many of the areas identified as having peat-forming potential are located outside existing protected areas, leaving them vulnerable to land-use changes, agriculture, and urban expansion (Fernandes et al., 2024). For example, the southwestern Iberian Peninsula and parts of northern Morocco exhibit moderate to high richness values but lack adequate protection, emphasizing the need for targeted conservation measures. For example, in Europe, only 20% of peatlands are located within protected areas (United Nations Environment Programme, 2024). Particularly, Portugal is one of the European countries with the highest proportion of currently degraded peatland area (Tanneberger, Moen, et al., 2021). Therefore, the effort to identify and map possible peatland areas that are not known is necessary to promote their conservation and protection. The richness maps generated in this study can serve as strategic tools for guiding field surveys, by identifying areas with high potential for peatland formation, even in fragmented or overlooked regions such as southern Portugal, Southwestern Spain, northern Morocco, northeastern Algeria and northern Tunisia. Furthermore, the vulnerability of peatlands to climate change, particularly in the WM (IPCC, 2023), underscores the urgency of incorporating these findings into climate-resilient conservation frameworks.

While this study provides valuable insights, it is not without limitations. The quality of occurrence data, particularly for less-studied regions like northern Morocco, may introduce biases due to uneven sampling effort or incomplete records (Campbell et al., 2021). Additionally, the reliance on environmental proxies, while effective, may not fully capture the ecological complexity of peat-forming and accumulating environments, particularly in regions with unique hydrological or climatic conditions. Complementary approaches, such as remote sensing and detailed field surveys, could be integrated to validate predicted richness areas and refine mapping efforts (Minasny et al., 2019). Furthermore, incorporating climate change scenarios into SDMs would provide valuable insights into the future distribution and resilience of peat-forming ecosystems in the WM.

## Conclusion

Our work highlights the effectiveness of bioindicator-based SDMs in predicting the distribution of peatland and organic soil-forming environments in the Western Mediterranean. The findings reveal significant areas of high richness, including overlooked regions such as southern Portugal and northern Morocco, northeastern Algeria, and northern Tunisia, providing a critical baseline for conservation planning. The methodological approach, combining cross-taxa richness modelling with a comprehensive set of environmental predictors, demonstrates the potential of SDMs for advancing peatland mapping in underrepresented regions. By supporting global efforts to map and conserve peatlands, this study contributes to the broader goal of protecting these critical ecosystems and the services they provide in the face of ongoing environmental change.

## CRediT authorship contribution statement

**Miguel Geraldes:** Conceptualization, Investigation, Methodology, Writing – original draft, Writing – review & editing. **Raquel Fernandes:** Conceptualization, Formal analysis, Investigation, Methodology, Writing – original draft, Writing – review & editing. **Maurício Santos:** Conceptualization, Methodology. **César Capinha:** Conceptualization, Methodology, Supervision, Writing – original draft, Writing – review & editing.

## Acknowledgements

RF was supported by a grant (PRT/BD/153505/2021) financed by the Portuguese Foundation for Science and Technology (FCT) under the MIT Portugal Program, Portugal. This research was supported by FCT, under the PEATMAP project grant (Ref. 2022.15748.MIT; http://doi.org/10.54499/2022.15748.MIT).

